# G2-to-G0 cell cycle exit underlies sensitivity to ATR inhibition via the p53-p21-RB1 axis

**DOI:** 10.64898/2026.04.28.721427

**Authors:** Celina Sanchez, Carlos Origel Marmolejo, Juyoung Lee, Joshua C. Saldivar

## Abstract

Ataxia-telangiectasia and Rad3-related (ATR) is an essential DNA damage response kinase that protects genome integrity by controlling cell cycle checkpoints, regulating origin firing, stabilizing replication forks, and signaling DNA repair. Due to hyper-proliferation, cancer cells depend on ATR for survival, implicating ATR inhibitors as promising therapeutics. However, variable tumor responses to ATR inhibitors highlights the need to uncover the determinants of cell fate. Here, we show breast cancer sensitivity to ATR inhibition correlates with the appearance of pan-nuclear DNA damage. The fate of these cells is driven by a p53-p21-RB1 axis that triggers a G2-to-G0-like cell cycle exit and is buffered by the p53 inhibitor MDM2. MDM2 inhibition lowers the DNA damage threshold for cell cycle exit and robustly targets ATR inhibitor-resistant cells. Our work reveals cell cycle plasticity as a mechanism determining cell fate during ATR inhibition and identifies MDM2 as a target for increasing ATR inhibitor efficacy.

## INTRODUCTION

The cell cycle is tightly regulated to ensure DNA replication and mitosis are successfully completed each round of cell division^1^. Its orderly progression begins with the decision to proliferate or enter a G0-like state at the end of mitosis^2^. Retinoblastoma 1 (RB1) is a key regulator of the G0-G1 transition^3^, acting to maintain quiescence of G0 cells when hypophosphorylated. Upon hyper-phosphorylation of RB1, cells enter G1 and, following inactivation of APC/C^Cdh1^, commit to S phase entry^4^. Progression through S phase is driven by replication origin firing, a process that resembles a specific timing program, coordinated by the activities of multiple cyclin-dependent kinases (CDKs) and associated cyclins, including cyclin E-CDK2, cyclin A-CDK2, and cyclin A-CDK1^5–7^. At the S/G2 transition, cyclin B levels begin to increase and promote the rise in mitotic CDK1-cyclin B activity^8^. Collectively, CDK1/2 activity increases during cell cycle progression, peaking at the end of the G2 phase to trigger mitotic entry.

Counteracting CDK1/2 is the ATR-CHK1 pathway^9^, which in conjunction with WEE1 kinase, functions to prevent hyperactivation of CDK1/2 in S and G2 phases. ATR-CHK1 signaling is critical for the orderly progression from S phase to mitosis^10^, as inhibition of the pathway deregulates the S/G2 transition, leading to premature expression of the G2/M gene network and mitotic entry with under-replicated chromosomes^11^. Consequently, cells acquire mitotic errors that are transmitted to daughter G0/G1 cells.

Mechanistically, ATR and its binding partner, ATRIP, are recruited to single-stranded DNA coated with RPA^12,13^. This platform arises frequently during DNA replication when replisomes encounter DNA damage that impedes replisome function^14^. Recruitment of the ATR activators, TOBP1 and/or ETAA1^15–18^, stimulates ATR kinase activity leading to CHK1 phosphorylation and activation^19^. CHK1 then phosphorylates the CDC25 phosphatases leading to their sequestration or degradation^20^. This then leads to CDK1/2 inhibition, as CDC25 phosphatases are needed to dephosphorylate inhibitory phosphorylations at threonine-14/tyrosine-15 of CDK1/2.

As ATR inhibition disrupts cell cycle checkpoints and impairs DNA repair, ATR inhibitors (ATRi) continue to be evaluated in clinical trials in a variety of aggressive cancer types, though their efficacy has been variable^21,22^. While biomarkers such as ATM and BRCA1/2 deficiency have been identified as predictors of response to ATRi, a more comprehensive understanding of the cellular mechanisms driving sensitivity to ATR inhibition is needed to improve therapeutic outcomes^23,24^.

Recently, we showed that ATR activity in S phase regulates the expression of the replication-dependent histones. This function of ATR occurs through local regulation of CDK1/2 activity within the histone locus body, a nuclear compartment that forms around histone gene clusters and couples histone expression to S phase. Loss of this function of ATR triggers over-expression of linker histones that then facilitate a nucleus-wide DNA damage signaling event visualized as pan-nuclear γH2AX staining^25^. Despite the dramatic presentation of this phenotype, the underlying signaling events and the eventual fate of these cells remain unknown.

Here, we utilize high-throughput quantitative image-based cytometry^26^ to examine the mechanism underlying pan-nuclear DNA damage, revealing that these cells go through a G2-to-G0 cell cycle exit via the p53-p21 pathway and rely on MDM2 for survival.

## RESULTS

### ATR inhibition drives distinct pathological cell fates

ATR regulates CDK1 and CDK2 activities in S phase, and in doing so impacts multiple processes in S phase, including origin firing, replication fork stability, S/G2 checkpoint signaling, transcription condensate dissolution, and histone gene regulation^9,11,25^. Yet the fate of cells subjected to ATR inhibition remains poorly defined. We treated MCF10A cells, a non-malignant epithelial cell line, with the ATR inhibitor, AZD6738, for 24 hours and pulsed with EdU for the last 15 minutes to label the S phase population. Following ATR inhibition, distinct cellular fates were identified, with most of the population exhibiting minimal DNA damage foci (Fig. 1 A). A fraction of cells showed γH2AX staining throughout the nucleus, which we recently referred to as pan-nuclear^25^. Micronuclei appeared in roughly 15 percent of cells, which can be caused by lagging chromosomes or chromosome fragmentation during mitosis^27–29^. Furthermore, a small percentage of cells underwent mitotic catastrophe, an event likely resulting from improper activation of cell cycle checkpoints, improper chromosomal segregation, and/or activation of apoptotic pathways during mitosis^30^ (Fig. 1 B). These diverse cell fates were partially linked to different cell cycle phases. For example, micronuclei were primarily observed in G1, while DNA damage foci were present at similar levels throughout the cell cycle. Interestingly, cells exhibited pan-nuclear damage from mid-S phase until G2 (Fig. 1 C). Given the cell cycle specific appearance of pan-nuclear γH2AX and its striking appearance, we chose to pursue further analysis of the fate of cells presenting this phenotype (Fig. 1 D).

**Figure 1.**
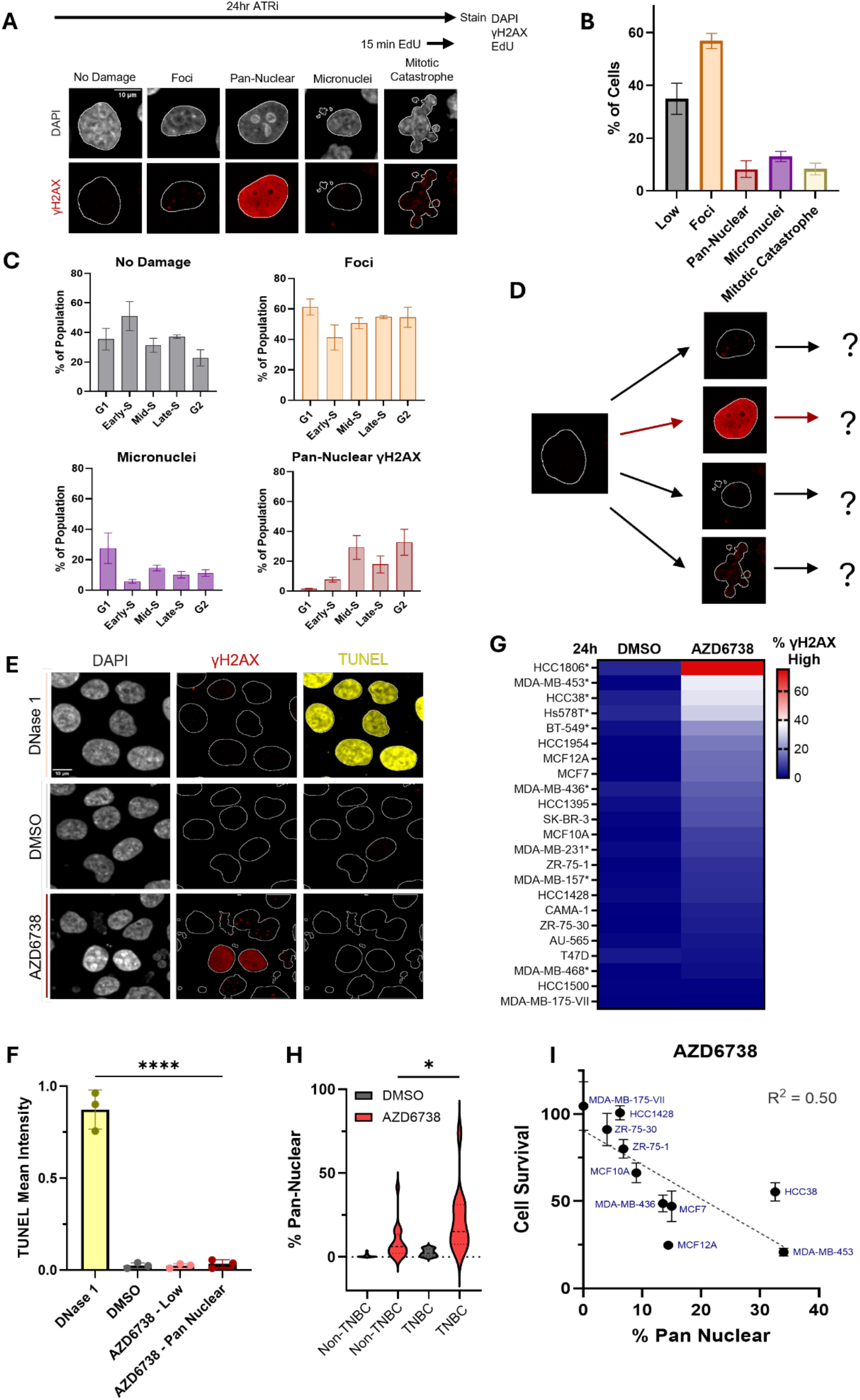
Pan-nuclear γH2AX formation correlates with ATRi sensitivity. A. Schematic indicating experimental setup and representative images of different DNA damage fates after 24 hours of ATR inhibition with 5 μM AZD6738. Scale bar, 10 μm. B. Bar graphs showing the percent of cells with the different DNA damage fates treated as in (A). C. Bar graphs of the percentage of cells with the different DNA damage fates in each phase of the cell cycle treated as in (A). D. Schematic of the various types of DNA damage fates that result from ATR inhibition. E. Representative images of γH2AX and TUNEL in cells treated as in (A). DNase I is a positive control for TUNEL. Scale bar, 10 μm. F. Bar graphs of the mean intensity of TUNEL. Cells were treated as in (A). AZD6738 cells were split based on γH2AX level (normal vs pan-nuclear). G. Heatmap of 21 breast cancer cell lines showing the percent of cells with pan-nuclear γH2AX treated as in (A). (*) indicates a triple negative breast cancer cell line (TNBC). H. Violin plots of the percentage of cells with pan-nuclear γH2AX in Non-TNBC versus TNBC. Cell lines were treated as in (A). I. Scatterplot of the percentage of cells with pan-nuclear γH2AX after 24 hours of treatment with 5 μM AZD6738 versus the cell survival relative to DMSO after 48 hours of treatment with 1 μM AZD6738 in ten breast cancer cell lines.

We first asked if the pan-nuclear γH2AX in ATR-inhibited cells was a result of apoptosis-induced DNA fragmentation, as high levels of γH2AX has been previously shown to co-occur with TUNEL-positive apoptotic staining after irradiation^31^. We used DNase I as a positive control for widespread DNA breaks in a TUNEL assay, as it is an endonuclease that cleaves DNA into fragments throughout the nucleus. However, during ATR inhibition alone, pan-nuclear γH2AX did not correlate with TUNEL staining, indicating these cells were not being primed for apoptosis nor were their nuclei full of fragmented DNA (Fig. 1, E and F).

### Pan-nuclear γH2AX formation correlates with cell survival and the TNBC subtype

Next, we asked if the pan-nuclear γH2AX correlated with cell survival and sensitivity to ATR inhibition. For this, we expanded our analysis to a panel of 21 breast cancer cell lines, with MCF10A cells serving as a non-malignant control. We observed variable accumulation of pan-nuclear γH2AX, with some cancer cell lines forming pan-nuclear staining more frequently than MCF10A cells and others less (Fig. 1 G). The cell lines were further characterized as triple negative breast cancer (TNBC) or non-triple negative breast cancer (Non-TNBC), except for HCC1500, since its subtype is contested^32^. The TNBC lines had a significantly higher percentage of cells with pan-nuclear γH2AX accumulation following ATR inhibition than the non-TNBC lines (Fig. 1 H), consistent with preclinical studies showing ATR inhibitors have a stronger therapeutic effect in TNBC^33^. To determine whether pan-nuclear γH2AX is a marker of long-term sensitivity to ATR inhibition, we examined how extended exposure affected survival in 11 of the breast cell lines, with MCF10A again used as a control. We observed a consistent decrease in cell survival as the population of γH2AX positive cells increased (Fig. 1 I). This suggests that the pan-nuclear phenotype is linked to the overall sensitivity to ATR inhibition and may serve as an informative biomarker for therapeutic efficacy of ATR inhibitors. Moreover, this highlights the importance to elucidate the underlying mechanism and fate of cells presenting pan-nuclear γH2AX.

### p53 supports DNA replication in cells with pan-nuclear DNA damage

p53 is a transcription factor that, in response to DNA damage, upregulates p21, a potent inhibitor of CDKs, leading to arrest in either G1 or G2^2,34^. To examine whether p53 influenced cell fate following ATR inhibition, we investigated the response to ATR inhibitors in p53^−/−^ MCF10A cells. First, we imaged p53 and quantified its levels and observed a striking upregulation of p53 following ATR inhibition. Importantly, p53 knockout cells did not express p53, confirming the loss of p53 in these cells (Fig. S1 A). We tested if p53 upregulation was specific to the population of cells with pan-nuclear γH2AX. As expected, we found that p53 was upregulated in this population of both MCF10A and the non-malignant retinal pigment epithelial (RPE-1 hTERT) cells (Fig. 2, A and B). Additionally, CHK1 inhibition also increased expression of the p53 pathway in cells with pan-nuclear γH2AX (Fig. S1, B).

**Figure 2.**
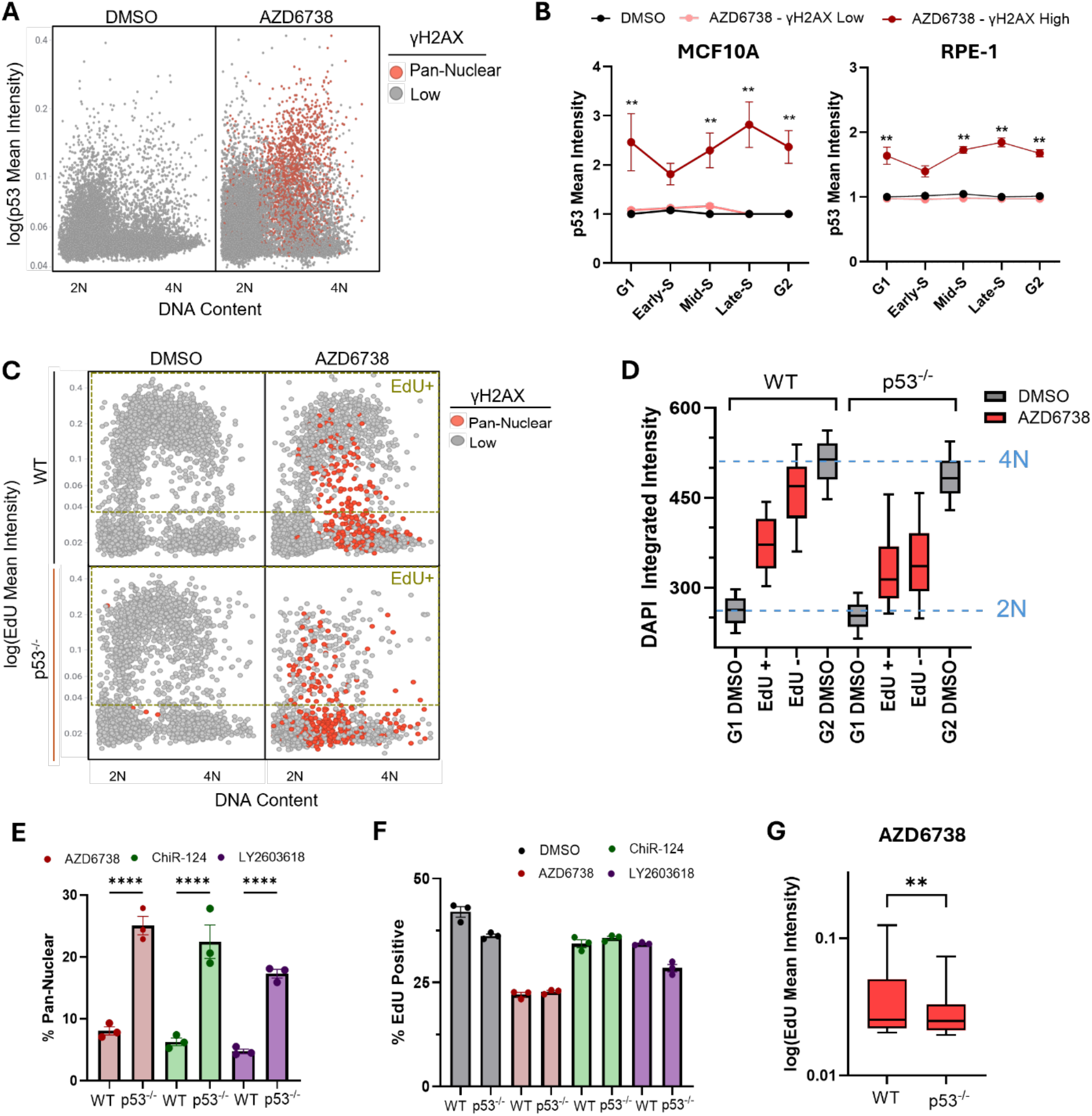
p53 is required for continued replication in cells with pan-nuclear γH2AX. A. Scatterplots of DNA content (DAPI integrated intensity) versus p53 mean Intensity (log_2_ scale) in MCF10A cells treated with DMSO (control) or 5 μM AZD6738 for 24 hours. Cells positive for pan-nuclear γH2AX were colored red. B. Line plots of p53 mean intensity in each cell cycle phase of MCF10A and RPE-1 cells. Cells were treated as in (A). Dark red line shows levels in pan-nuclear γH2AX (γH2AX High) cells. C. Scatterplots of DNA content (DAPI integrated intensity) versus EdU mean intensity (log_2_ scale) in MCF10A and p53^−/−^ MCF10A cells treated with or 5 μM AZD6738 for 24 hours. Cells that were EdU positive are indicated by the yellow box. Cells positive for pan-nuclear γH2AX were colored red. D. Boxplots of the DAPI integrated intensity in MCF10A and p53^−/−^ MCF10A cells with pan nuclear γH2AX (red-shaded boxes) following 24 hours of treatment with 5 μM AZD6738. Gray-shaded boxes show the DAPI intensity in DMSO-treated G1 and G2 cells. G1 and G2 cells were used to mark the 2n and 4n DNA content, respectively. Boxes and whiskers show the 25-75^th^ percentile and 10-90^th^ percentile, respectively. E. Bar graphs of the percentage of cells with pan-nuclear DNA damage during AZD6738 (red, 5 μM), ChiR-124 (green, 250 nM), or LY2603618 (purple, 2 μM) treatment for 24 hours. F. Bar graphs of the percentage of EdU positive cells. Cells were treated as in (C). G. Boxplots of EdU mean intensities in WT-MCF10A and p53^−/−^ MCF10A cells with pan-nuclear DNA damage during treatment with AZD6738 (5 μM) for 24 hours. Boxes and whiskers show the 25-75^th^ percentile and 10-90^th^ percentile, respectively.

Next, we examined the impact of p53 loss on the response to ATR inhibition. Within 2 hours of ATR inhibition, the p53^−/−^ cells showed similar accumulation of pan-nuclear cells in mid-S phase; however, the impact of p53 loss became apparent after prolonged inhibition (Fig. S1, C-E) Specifically, p53-WT cells continued replicating in the presence of pan-nuclear γH2AX and eventually exited S phase with close to 4N DNA content. However, the p53^−/−^ cells were unable to replicate with pan-nuclear γH2AX and exited S phase with at most 3N DNA content (Fig. 2, C and D). Similarly, inhibition of the downstream target of ATR, CHK1, also resulted in a failure to replicate in p53^−/−^ cells with pan-nuclear DNA damage (Fig. S1, F). Furthermore, p53 loss led to a higher percentage of the population presenting with pan-nuclear DNA damage during ATR and CHK1 inhibition (Fig. 2 E), which is consistent with other reports showing increased DNA damage in p53-deficient cells^35^. Additionally, the majority of the p53^−/−^ cell population with pan-nuclear DNA damage stopped replicating, while the overall population of pan-nuclear cells in control MCF10A cells was still capable of incorporating EdU, albeit at a lower efficiency compared to undamaged cells (Fig. 2, F and G). Overall, these data suggests that p53 is required for cells with pan-nuclear γH2AX to finish replicating in S phase and is a driver of cell fate during ATR inhibition.

### The p53-p21 pathway induces a G2-to-G0 cell cycle exit in response to ATR inhibition

Activation of p53 leads to transcriptional activation of p21, which binds to CDKs and inhibits their kinase activities, to induce cell cycle arrest^36,37^. Accordingly, we examined the levels of p21 following ATR-CHK1 inhibition. Notably, there was a significant increase in p21 levels across the cell cycle in pan-nuclear cells during ATR and CHK1 inhibition (Fig. 3, A and B, and Fig. S2 A). Next, we used an established DNA-helicase B (DHB)-based CDK2 biosensor^2^ to determine if p21 upregulation and pan-nuclear γH2AX correlated with a loss of CDK activity. The biosensor utilizes DHB translocation to track CDK2 activity across the phase of the cell cycle (Fig. 3 C). MCF10A-DHB cells exhibited increased CDK2 activity in the 4N population compared to 2N; however, the cells with pan-nuclear DNA damage had reduced CDK2 activity in the 4N population, consistent with the rise in p21 levels acting to inhibit CDK1/2 (Fig. 3, D and E). Furthermore, single-cell traces of CDK2 activity from time-lapse images of cells treated for 24 h with ATRi revealed an initial rise in CDK2 activity consistent with S-G2 progression, but then a sharp decline in CDK2 activity in cells with pan-nuclear DNA damage to levels similar to G0-cells despite not entering and transversing mitosis (Fig. S2 B). This data suggests that damaged cells enter G2, but do not divide, and instead turn off CDK2 activity and exit the cell cycle.

**Figure 3.**
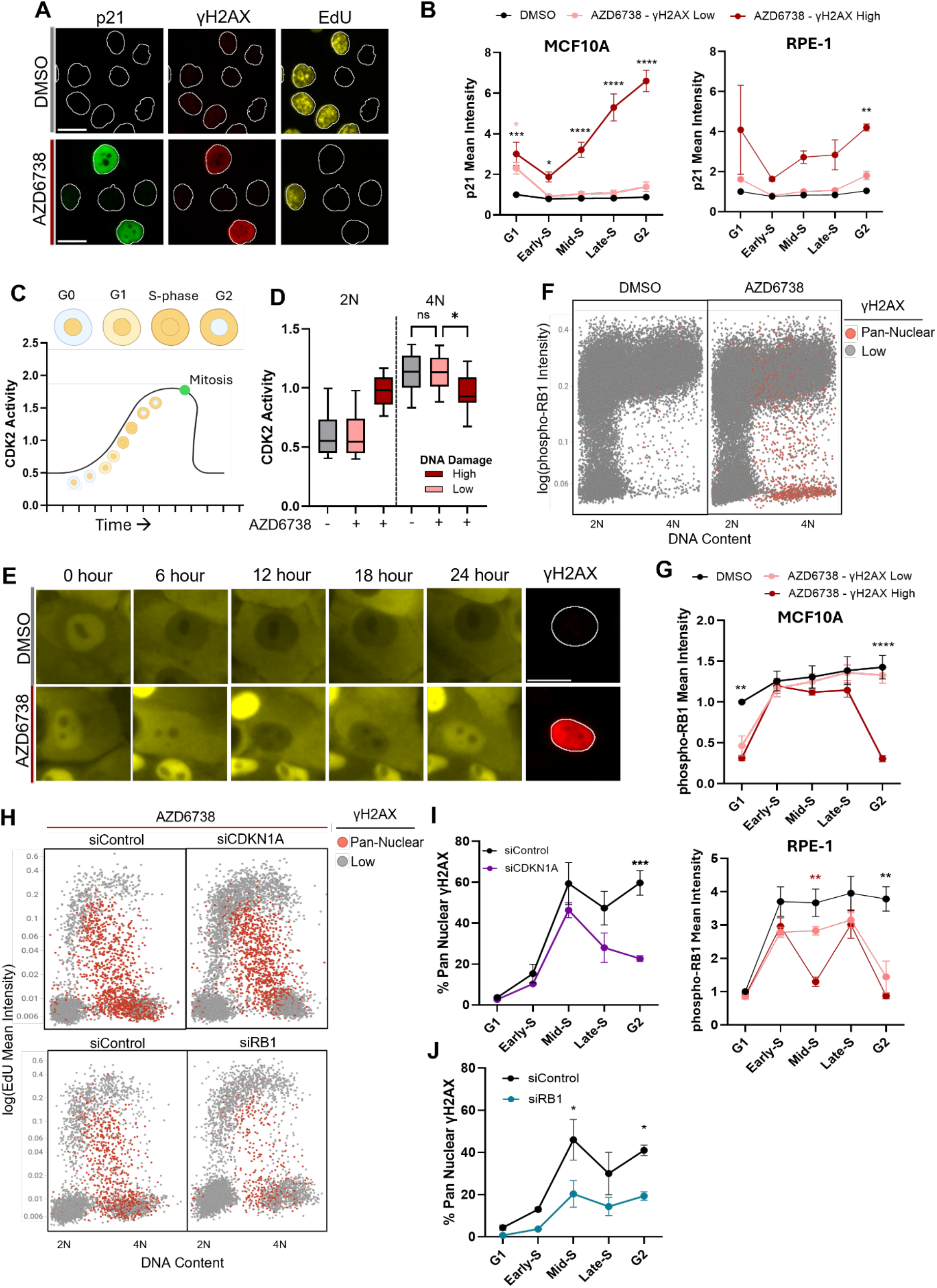
ATR inhibition promotes a G2-to-G0 cell cycle exit. A. Representative images of p21, γH2AX, and EdU in MCF10A cells. Cells were treated with DMSO or 5 μM AZD6738 for 24 hours. Scale bar, 10 μm. B. Line plots of p21 mean intensity in each cell cycle phase of MCF10A and RPE-1 cells. Cells were treated as in (A). Dark red line shows levels in pan-nuclear γH2AX (γH2AX High) cells. C. Schematic of how CDK2 phosphorylation activity causes translocation of DHB-mVENUS from the nucleus to the cytoplasm at different stages of the cell cycle. D. Boxplots of CDK2 activity in cells with 2N and 4N DNA content. Cells were treated with DMSO or 5 μM AZD6738 for 24 hours. Dark red box shows levels in pan-nuclear γH2AX (γH2AX High) cells. E. Representative images of DHB-mVENUS at 6, 12, 18, and 24 hours and γH2AX after 24 hours of treatment described in (D). Scale bar, 10 μm. F. Scatterplots of DNA content (DAPI integrated intensity) versus phospho-RB1 mean intensity (log_2_ scale) in MCF10A cells treated as in (A). Cells positive for pan-nuclear γH2AX were colored red. G. Line plots of phospho-RB1 mean intensity across the cell cycle in MCF10A and RPE-1 cells. Cells were treated as in (A). Dark red line shows levels in pan-nuclear γH2AX (γH2AX High) cells. Boxes and whiskers show the 25-75^th^ percentile and 10-90^th^ percentile, respectively. H. Scatterplots of DNA content (DAPI integrated intensity) versus EdU mean intensity (log_2_ scale) in MCF10A cells transfected with non-targeting siRNA (siControl), siCDKN1A (p21), and siRB1 and treated with 5 μM AZD6738 for 24 hours. Cells positive for pan-nuclear γH2AX were colored red. I. Line plots of the percentage of pan-nuclear γH2AX cells across the cell cycle after siRNA transfection with siControl and siCDKN1A and treated with 5 μM AZD6738 for 24 hours. J. Line plots of the percentage of pan-nuclear γH2AX cells across the cell cycle after siRNA transfection with siControl and siRB1 and treated as in (I).

Recently, it was demonstrated that loss of CDK4/6 activity in G2 could lead to retinoblastoma protein (RB1) dephosphorylation, suggesting the regulation of RB1 via phosphorylation could extend beyond the G0/G1 restriction point^38,39^. Classically, hypophosphorylated RB1 binds and inhibits E2F transcription factors and thereby arrests cells in G0. When RB1 is phosphorylated by CDK4/6, the inhibition of E2F is relieved, allowing bypass of the restriction point and cell cycle commitment. Consistent with this model, phosphorylation of RB1 appeared switch-like in that 2N cells were clearly divided between the hypophosphorylated state (G0 cells) and the hyperphosphorylated state (G1 cells) (Fig. 3 F). RB1 hyperphosphorylation persisted as cells progressed through S and G2 phases. Strikingly, following ATR inhibition, there was a switch back to RB1 hypophosphorylation in the G2 phase, specifically in cells with pan-nuclear γH2AX (Fig. 3, F and G). CHK1 inhibited cells also reactivated RB1 in the G2 phase and entered a G0-like state (Fig. S2 C). To determine if RB1 was inactivated or degraded, we co-stained for total RB1 and γH2AX. Interestingly, there was a partial, yet significant, decrease in total RB1 in G2 pan-nuclear cells during ATR and CHK1 inhibition, suggesting RB1 is targeted for degradation after being inactivated (Fig. S2, D-F).

Since p53 is critical for damaged cells to finish replicating under ATR inhibition, we examined if the p53 pathway was dysfunctional in p53^−/−^ MCF10A cells. Somewhat unexpectedly, p21 was moderately elevated in G1 cells compared to the rest of the cell cycle phases despite the absence of p53 (Fig. S2 G). Of note, ATR-inhibited cells with pan-nuclear DNA damage did not upregulate p21 in the p53-null cells, consistent with p21 acting downstream of p53 following ATR inhibition (Fig. S2 G). RB1 phosphorylation in p53^−/−^ cells was similar to p53-WT cells; however, during ATR inhibition, knockout cells experienced hypophosphorylation of RB1 in G1 and G2 phases, independent of DNA damage (Fig. S2 H).

Next, we examined the effect of p21 and RB1 knockdown (KD) on cells with pan-nuclear DNA damage (Fig. S3 A). Notably, there was no difference in the percentage of cells in each phase of the cell cycle following KD (Fig. S3 B). However, both p21 and RB1 KD decreased the percent of the population with pan-nuclear γH2AX (Fig. S3 C), albeit with differing cell cycle specificities. For p21 KD, cells accumulated pan-nuclear γH2AX in mid- and late-S phase, but in G2 the damage was not present, suggesting p21 acts to sustain the DNA damage marker in G2 cells (Fig. 3, H and I). RB1 was highly phosphorylated in p21 KD cells throughout the entirety of the cell cycle, which may contribute to the abrupt switch in DNA damage signaling (Fig. S3 D). RB1 knockdown prevented the accumulation of pan-nuclear γH2AX in mid-S phase through G2, suggesting RB1 promotes the spreading of this DDR marker throughout the nucleus (Fig. 3 J). Altogether, the data show that the components of the cell cycle exit pathway play a critical role in the development and continuous signaling of the DDR to generate the pan-nuclear γH2AX phenotype.

### ATR inhibition induces MDM2 expression

To further explore the molecular pathways regulating cell fate in response to ATR inhibition, we measured active gene transcription in ATR-inhibited cells via nascent RNA-sequencing. MCF10A cells were treated with DMSO or AZD6738 for 8 hours prior to labeling of nascent RNA by EU incorporation. EU-labeled RNA was biotinylated via click chemistry, affinity-purified, converted to cDNA, and subjected to sequencing (Fig. 4 A). The top two upregulated genes in ATR-inhibited cells were *CDKN1A* and *MDM2*, both transcriptional targets of p53 (Fig. 4 B). *CDKN1A* encodes for p21 and upregulation of this gene, correlates with the observed increase in p21 in cells with pan-nuclear DNA damage (Fig. 3 B). Gene ontology analysis for the top 25 upregulated genes showed an increase in apoptotic signaling pathway genes and the DNA damage response, suggesting that during ATR inhibition cells are amassing a p53-dependent transcriptional response to significant levels of DNA damage (Fig. 4 C).

**Figure 4.**
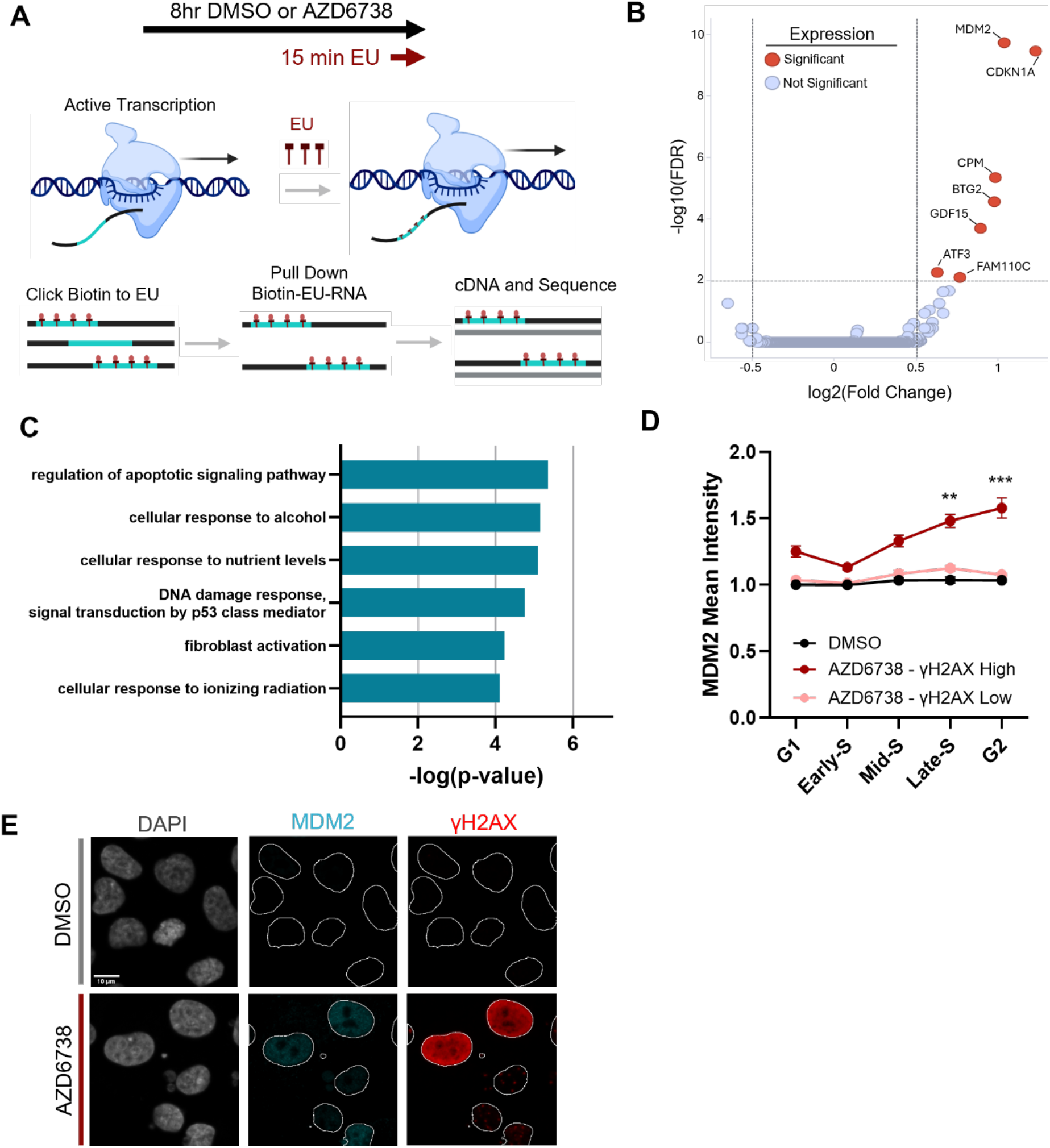
ATR inhibition triggers MDM2 upregulation. A. Schematic of experimental setup and sample preparation for nascent RNA-sequencing. B. Volcano plot of differentially expressed genes in MCF10A cells treated with 5 μM AZD6738 for 8 hours. Fold change was relative to DMSO-treated MCF10A cells. Genes that are significant upregulated with ATRi are labeled red. C. Gene ontology analysis of the top 25 differentially expressed genes from (B). D. Line plots of MDM2 mean intensity across the cell cycle of MCF10A cells treated with DMSO or 5 μM AZD6738 for 24 hours. Dark red line shows levels in pan-nuclear γH2AX (γH2AX High) cells. E. Representative images of DAPI, MDM2, and γH2AX in MCF10A cells. Cells were treated as in (D). Scale bar, 10 μm.

MDM2 (murine double mutant 2) is a negative regulator of p53 and has oncogenic activity in various cancer contexts. Given that it was among the most highly upregulated genes with ATR inhibition, we assessed its functional role in ATRi and CHK1i-induced cells with pan-nuclear DNA damage. Like p21, MDM2 levels were elevated in late-S and G2 cells with pan-nuclear γH2AX staining (Fig. 4, D and E, and Fig. S4 A). Knockdown of MDM2 (Fig. S4, B and C) did not significantly affect the accumulation of γH2AX despite it being upregulated, suggesting its role as a p53 inhibitor may influence the G2-to-G0 cell cycle exit decision without altering DNA damage or DDR signaling.

MDM2 is a candidate target in cancer therapy, given its oncogenic role as an inhibitor of p53 tumor suppressor activity. Numerous inhibitors of MDM2 (MDM2i) have been developed, showing that restoration of p53 signaling can potentiate p53-mediated apoptosis^40–42^. MDM2i are currently being evaluated in clinical trials as both a monotherapy and in combination with other drugs^43,44^. We tested two MDM2i, idasanutlin and nutlin-3a, to determine if the combination with AZD6738 would increase p21 activation and alter cell fate. We noted a marked decrease in the percentage of cells in S-phase during dual inhibition for 24 hours (Fig. S4 D). The combination of MDM2i and ATRi did not impact DNA damage accumulation; however, the combination increased p21 levels in both the G1 and G2 phases compared to either inhibitor alone (Fig. 5 A and Fig. S4, E and F). Particularly, the combination of MDM2i and ATRi, caused nearly all G2 cells, independent of pan-nuclear DNA damage, to dephosphorylate RB1 and undergo a G2-to-G0 exit (Fig. 5, B and C, and Fig. S4, G and H). Similarly, we observed a loss of total RB in the entire G2 population during the combination treatment, again, irrespective of pan-nuclear γH2AX presentation (Fig. S4 I). We also observed an increase in the size of the nuclei of G2 cells, indicative of a senescent-like phenotype, which was not as prominent in the other phases of the cell cycle (Fig. S4 J). Finally, we examined MDM2 inhibition and the effect on CDK2 activity using the CDK2 biosensor. For both MDM2 inhibition alone and in combination, we observed a loss of CDK2 activity in the 4N population independent of DNA damage (Fig. 5 D).

**Figure 5.**
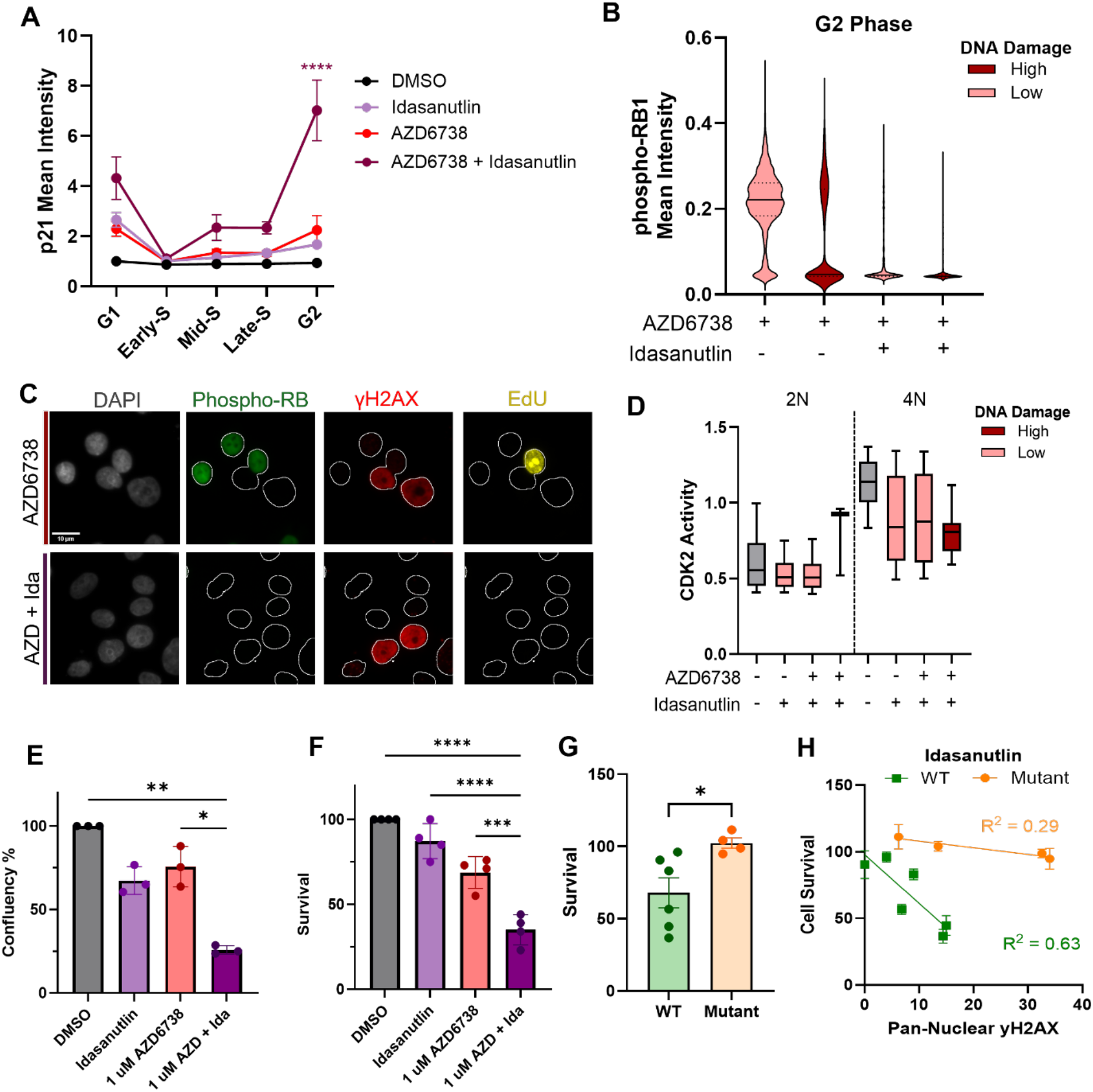
Combination of MDM2 and ATR inhibitors synergistically prevent cell proliferation. A. Line plots of p21 mean intensity across the cell cycle of MCF10A cells treated with DMSO, 50 nM idasanutlin, 5 μM AZD6738, or both for 24 hours. B. Violin plots of phospho-RB1 intensity in the G2 phase in MCF10A cells treated as in (A). Dark red plots show levels in pan-nuclear γH2AX (γH2AX High) cells. C. Representative images of DAPI, phospho-RB, γH2AX, and EdU. Cells were treated as in (A). Scale bar, 10 μm. D. Boxplots of CDK2 activity in cells with 2N and 4N DNA content. Cells were treated with DMSO, idasanutlin (50 nM), or the combination of idasanutlin (50 nM) and AZD6738 (5 μM) for 24 hours. Dark red color shows levels in pan-nuclear γH2AX (γH2AX High) cells. Boxes and whiskers show the 25-75^th^ percentile and 10-90^th^ percentile, respectively. E. Relative cell number after treatment with DMSO, 50 nM idasanutlin, 1 μM AZD6738, or the combination for 48 hours. F. Cell viability of MCF10A cells treated as in (E) after 3 days of recovery with normal media. Viability was normalized to DMSO. G. Bar graphs of cell survival after 48 hour treatment with 50 nM idasanutlin and recovery across six WT-p53 and four mutant p53 breast cancer cell lines. H. Scatterplot of the percentage of pan-nuclear cells after 24 hours of treatment with AZD6738 (5 μM) versus the relative cell survival after 48 hours of treatment with idasanutlin (50 nM) across six WT-p53 and four mutant p53 breast cancer lines.

Next, we examined the effect of ATRi and MDM2i combination on cell viability. For this, we first performed dose-response viability curves on cells treated with individual inhibitors (Fig. S4 K), identifying concentrations that caused an approximate 25% reduction in viability. We then tested the effect of combining ATR and MDM2 inhibitors at these lower concentrations and observed a synergistic loss of viability compared to each inhibitor alone after 48 hours of treatment, suggesting ATR and MDM2 combined inhibition may be an effective anti-cancer strategy (Fig. 5, E and F).

To determine if the effect of the MDM2i was dependent on p53 status, we analyzed MDM2i response in six p53 WT and four p53 mutant breast cancer cell lines. As expected, MDM2 inhibition decreased cell survival in half of the p53 WT cell lines but did not impact cell survival in mutant p53 cell lines (Fig. 5 G). Interestingly, sensitivity to MDM2 inhibition in the p53 WT cell lines correlated with the percentage of cells with pan-nuclear DNA damage during ATR inhibition (Fig. 5 H). This relationship suggests a link between ATR and MDM2 that underlies the sensitivity to these inhibitors. Altogether, our data show that the p53-p21 pathway is active in ATR-inhibited cells, but MDM2 suppresses G2-to-G0 cell cycle exit in cells with DNA damage levels below the pan-nuclear threshold. Importantly, these findings suggest the combination of MDM2 and ATR inhibitors may yield effective therapeutic responses in cancers with wild-type p53 or MDM2 amplification.

### MDM2 mediates resistance to ATR inhibition

A current challenge in utilizing ATR inhibitors as a cancer therapeutic is the potential for cancer cells to develop resistance. In clinical trials where ATR inhibitors are utilized as a monotherapy, treatment plans are often cyclic with 3-5 days on and 2 days off^24^. This type of treatment plan allows for a subpopulation of cells to develop resistance and promote relapse. Loss of genes such as *UPF2, CCNC*, and *CDK8* have been previously identified as mechanisms of resistance to ATR inhibitors^45,46^. However, little information is known about how cell cycle plasticity impacts cell fate and ultimately resistance to ATR inhibition. Thus, we utilized ATR inhibitor-resistant MCF10A cells (MCF10A-R) that we developed via long-term growth of cells in gradually-increasing sub-lethal doses of AZD6738. In stark contrast to MCF10A cells, MCF10A-R cells do not have pan-nuclear DNA damage and do not activate p21 during S-phase (Fig. 6, A and B). However, these cells exhibit a highly active G1 checkpoint that prevents or slows entry into S-phase (Fig. 6 C).

**Figure 6.**
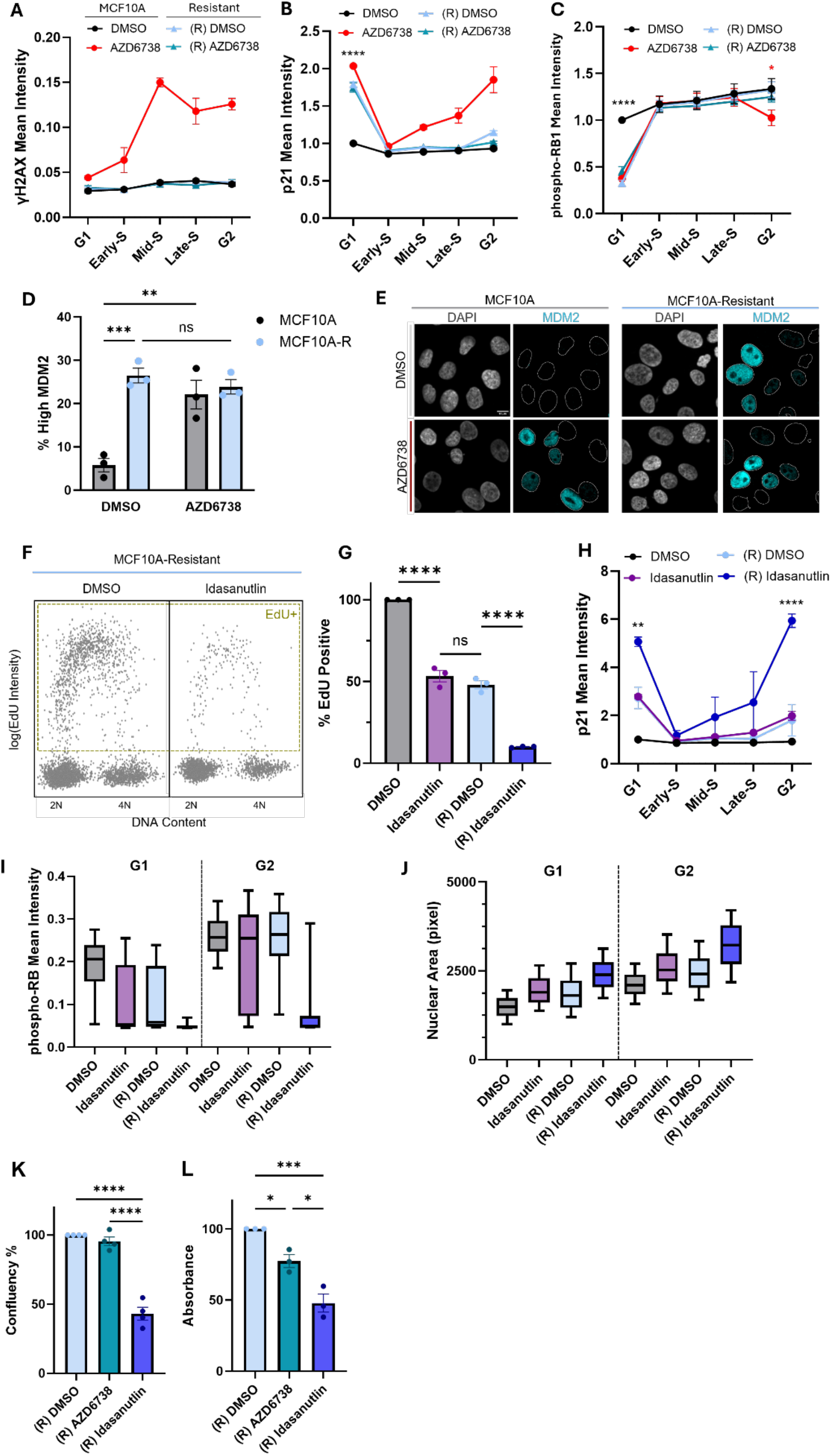
MDM2 overexpression promotes ATR inhibitor resistance. A. Line plots of γH2AX mean intensity across the cell cycle in parental MCF10A and MCF10A-Resistant (R) cells treated with DMSO or 5 μM AZD6738 for 24 hours. B. Line plots of p21 mean intensity across the cell cycle in cells treated as in (A). C. Line plots of phospho-RB mean intensity across the cell cycle in cells treated as in (A). D. Bar graphs of the percentage of MDM2 overexpression in parental MCF10A and MCF10A-Resistant cells. Cells were treated as in (A). E. Representative images of DAPI and MDM2 in parental MCF10A and MCF10A-Resistant cells. Cells were treated as in (A). F. Scatterplots of DNA content (DAPI integrated intensity) vs EdU Mean Intensity (log_2_ scale) in MCF10A-Resistant cells treated with DMSO or 50 nM Idasanutlin. Cells that were EdU positive are grouped in yellow box. G. Bar graphs of the percentage of EdU positive cells in parental MCF10A and MCF10A-Resistant cells during treatment with DMSO or 50 nM idasanutlin for 24 hours. Percentages were normalized to parental cells treated with DMSO. H. Line plots of p21 mean intensity across the cell cycle in cells treated as in (G). I. Boxplots of phospho-RB1 mean intensity in G1 and G2 phases. Cells were treated as in (G). Boxes and whiskers show the 25-75^th^ percentile and 10-90^th^ percentile, respectively. J. Boxplots of nuclear area (pixels squared) in G1 and G2 phases. Cells were treated as in (G). Boxes and whiskers show the 25-75^th^ percentile and 10-90^th^ percentile, respectively. K. Bar graphs of MCF10A-Resistant cell counts relative to DMSO after treatment with DMSO, 1 μM AZD6738, or 50 nM idasanutlin for 48 hours. L. Bar graphs of cell viability of MCF10A-Resistant cells treated as in (K) after 3 days of recovery with normal media.

Next, we asked if MDM2 levels were altered in ATRi-resistant cells. Unlike MCF10A cells, where MDM2 is only activated in response to ATR inhibition and pan-nuclear DNA damage formation, MCF10A-R cells have elevated MDM2 activity during normal proliferation (Fig. 6, D and E). These data suggest that ATR inhibitor-resistant cells may be dependent on a balance of G0/G1 checkpoint activity and elevated levels of MDM2 for survival.

MDM2i have been shown to combat CDK4/6 inhibitor resistance by stabilizing p53 and activating p21^47^. Therefore, we tested if MDM2i could overcome resistance to ATRi. Treatment with idasanutlin in MCF10A-R cells led to a marked reduction in replicating cells compared to control (Fig. 6, F and G). Additionally, p21 levels in idasanutlin treated MCF10A-R cells were increased significantly in both G1 and G2 phases when compared to control MCF10A-R and normal MCF10A cells treated with idasanutlin alone (Fig. 6 H). Similar to the combined treatment in parental MCF10A cells, individual treatment with idasanutlin in MCF10A-R promoted hypophosphorylation of RB1 and increased the nuclear area for all G1 and G2 cells (Fig. 6, I and J). Finally, after 48 hours of MDM2 inhibition, cell survival was reduced by 50% (Fig. 6, K and L). Altogether, these data suggest MDM2 inhibition may be an effective strategy for targeting ATRi-resistant cells via p21 overexpression and a robust G2-to-G0 cell cycle exit.

## DISCUSSION

Our study uncovers the cell fate of ATR-inhibited cells that present with pan-nuclear DNA damage in the S phase, revealing a G2-to-G0 cell cycle exit driven by the p53-p21-RB1 axis and buffered by MDM2. Specifically, ATR inhibition upregulates the p53 pathway in S phase with high levels of DNA damage. Despite this damage and p53-p21 accumulation, these cells progress to the G2 phase, where they shut off CDK1/2 activity, dephosphorylate RB1, and enter a G0-like state. However, p53 activity also increases MDM2 levels, which antagonizes p53-p21 effectively raising the DNA damage threshold needed for the G2-to-G0 cell cycle exit.

There are several curious aspects to the cell cycle plasticity induced by ATR inhibition. First, p53 and p21 upregulation begin in S phase and are needed for progression until the G2 phase; however, RB1 dephosphorylation only occurs in G2 phase. The difference in cell cycle phase for these processes suggest DNA replication prevents RB1 dephosphorylation despite high p21 levels. Second, it is unclear how S phase cells with pan-nuclear DNA damage and p53/p21 upregulation continue to replicate given p21 should inhibit CDK2 and arrest the cells in S phase.

Intriguingly, p21 activity is not needed for the initial formation of pan-nuclear γH2AX in S phase, yet it is needed for the persistence of this signal into the G2 phase. We speculate that pan-nuclear γH2AX is reversible in G2 cells, perhaps via a phosphatase that is more active in G2 cells in a CDK-dependent manner. The rise in p21 as cells reach G2 would then trigger a loss of CDK activity and thus a decrease in phosphatase activity. Consistent with this hypothesis, time-lapse microscopy revealed an initial rise and peak in CDK2 activity followed by a rapid decline in kinase activity to G0 levels in cells that present the pan-nuclear γH2AX phenotype.

Importantly, the pan-nuclear DNA damage induced by ATR inhibition is a potential determinant of breast cancer response to ATR inhibition and is more commonly observed in TNBCs, an aggressive and difficult to treat subtype of breast cancer^48^. Accordingly, γH2AX pan-nuclear staining may be a suitable biomarker for ATRi response in the clinic. However, TNBCs frequently harbor TP53 mutations^49,50^, and how these mutations impact TNBC cell fate in response to ATR inhibition needs further investigation.

A major challenge to the use of targeted therapies is the development of therapy-resistant cancer cells. To determine if ATR inhibitor resistance is linked to the G2-to-G0 cell fate, we developed resistant MCF10A cells and found that they rely on overexpression of MDM2 for proliferation. We propose that MDM2 inhibition may be an effective approach to target cancer cells that develop resistance to ATR inhibitors. In addition, the combination of ATR and MDM2 inhibitors may synergistically eliminate cancer cells, allowing the use of these drugs at lower concentrations. This would be a potential breakthrough as the drug dosages typically needed to elicit tumor responses are associated with adverse side effects, such as neutropenia and anemia, that can lead to treatment discontinuation^22,51^.

ATR inhibitors continue to be tested in clinical studies as anti-cancer agents. Our work links the p53-p21-RB1 axis to the fate of cells treated with these pharmacological drugs, pointing to cell cycle plasticity involving a G2-to-G0 exit, buffered by MDM2, as a key determinant. p53 and RB1 are among the most frequently altered tumor suppressors in cancer, and in some cancers both are mutated. Thus, there is need for clear and consistent biomarkers for ATR inhibitor response, and pan-nuclear γH2AX staining is a promising candidate.

## METHODS

### Cell Lines

All cell lines were purchased from ATCC. MCF10A, p53^−/−^ MCF10A, and MCF12A cells were cultured in DMEM:F12 (Thermo Fisher Scientific, 11320082) base medium supplemented with 5% horse serum (Thermo Fisher Scientific, 16050122), 20ng/ml EGF (Peprotech, AF-100-15), 10 ug/ml insulin (Millipore Sigma, I-1882), 0.5 mg/ml hydrocortisone (Millipore Sigma, H-0888), 100 ng/ml cholera toxin (Millipore Sigma, C-8052), and 1x penicillin/streptomycin (Thermo Fisher Scientific, 15140122). MCF10A-resistant cells were cultured in MCF10A media with the addition of 5 uM AZD6738. MCF7, MDA-MB-175-VII, T47D, and MDA-MB-231 were cultured in DMEM:F12 (Thermo Fisher Scientific, 11320082) base medium supplemented with 10% FBS. AU-565, BT-549, HCC1428, ZR-75-1, ZR-75-30, HCC1395, HCC1806, HCC1954, HCC38, and HCC1500 were cultured in RPMI-1640 (Thermo Fisher Scientific, 11875093) base medium supplemented with 10% FBS. MDA-MB-453, MDA-MB-436, MDA-MB-157, Hs578T, CAMA-1, SK-BR-3, and MDA-MB-468 were cultured in DMEM (Thermo Fisher Scientific, 12320032) base medium supplemented with 10% FBS. RPE-1 cells were cultured in DMEM:F12 (Thermo Fisher Scientific, 11320082) base medium supplemented with 10% FBS and 0.01 mg/mL hygromycin B (Millipore Sigma, 10843555001).

### Antibodies

The following antibodies were used: pRb Ser807/Ser811 (D20B12, Cell Signaling Technology, no. 8516, 1:1000), Rb (4H1, Cell Signaling Technology, no. 9309, 1:1000), p53 (7F5, Cell Signaling Technology, no. 2527, 1:500), p21 Waf1/Cip1 (12D1, Cell Signaling Technology, no. 2947, 1:1000), phospho-Histone H2A.X Ser139 (20E3, Cell Signaling Technology, no. 9718, 1:500), and MDM2 (D1V2Z, Cell Signaling Technology, no. 86934, 1:500). The following secondary antibodies were used to visualize non-conjugated primary antibodies: goat anti-rabbit, Alexa Fluor 488 (Thermo Fisher Scientific, A11034, 1:1000) and goat anti-mouse, Alexa Fluor 488 (Thermo Fisher Scientific, A11034, 1:1000).

### Chemicals and Reagents

DAPI (D9542) was used at 5 ug/mL to stain the nuclei. The following small molecule inhibitors were used: ATRi (AZD6738 5 M, Selleck Chemicals, S7693), CHK1i (LY2603618 2 uM, Selleck Chemicals, S2626) and (ChiR-124 250 nM, Selleck Chemicals, S2683), MDM2i (Idasanutlin 50 nM, Selleck Chemicals, S7205) and (Nutlin-3a 1 μM, Selleck Chemicals, S8059). Stock concentrations of all drugs were made with DMSO.

### TUNEL Assay

MCF10A cells were seeded at 13,000 cell per well. After 24 hours, media was replaced with media containing DMSO (control) or an ATR inhibitor. After 24 hours, cells were fixed for 10 minutes with 4% para-formaldehyde. Following methanol permeabilization, 1 unit of DNase (Thermo Fisher Scientific, 18068015) was added to the respective wells and incubated for 30 minutes at room temperature. The Click-IT TUNEL assay 594 (Thermo Fisher Scientific, C10618) was utilized according to the manufacturer’s instructions.

### siRNA transfection

The following siRNA were used: ON-TARGETplus Human Non-targeting Control Pool (Horizon Discoveries, D-001810-10-05, 50nM), ON-TARGETplus Human RB1 SMARTpool (5925, Horizon Discoveries, L-003296-02-0005, 50nM), ON-TARGETplus Human CDKN1A SMARTpool (1026, Horizon Discoveries, L-003471-00-0005, 50nM), and ON-TARGETplus Human MDM2 SMARTpool (4193, Horizon Discoveries, L-003279-00-0005, 50nM). siRNAs were combined with DharmaFECT 1 (Horizon Discovery, T-2005-01) according to the manufacturer’s guidelines. Cells were seeded at 7,000 cells per well and siRNA was added to each well. 16 hours after reverse-transfection, media was replaced with normal growth media. After 8 hours of incubation with the normal growth media, media was replaced with media containing DMSO (control), ATR inhibitors, or MDM2 inhibitors at the desired concentrations.

### Immunofluorescence staining

Cells were seeded in glass-bottom 96-well plates (Cellvis P96-1.5P) such that cell density remained sub-confluent (80-90% confluent) until the end of the treatment. 15 minutes before fixation 10 uM 5-Ethynyl-2’-deoxyuridine (EdU) (Millipore Sigma, 900584) was added to the media. Cells were fixed with 4% paraformaldehyde diluted in phosphate-buffered saline (PBS) for 10 min and then permeabilized with ice-cold methanol for 10 minutes. The cells were washed with 1X PBS and blocked in 3% bovine serum albumin (BSA) diluted in 1X PBS for 5 min at room temperature. EdU was fluorescently labelled with the Click-iT Cell Reaction Buffer kit (Thermo Fisher Scientific, C10269) and Alexa Fluor™ 647 Azide (Thermo Fisher Scientific, A10277) according to the manufacturer’s guidelines. The cells were incubated with the Click-iT reaction for 30 minutes at room temperature on a bench rocker. The cells were then washed with 1X PBS, washed in 3% BSA, and blocked with 1% BSA for 30 min at room temperature on a bench rocker. Cells were then incubated overnight at 4℃ with primary antibodies diluted in 1% BSA. Cells were then washed 3 times with 1X PBS for 5 minutes on a bench rocker. Secondary antibodies and DAPI (Millipore Sigma, D9542, 5 ug/mL) for 1 hour at room temperature on a bench rocker. Cells were washed 3 times with 1X PBS for 5 minutes on a bench rocker before imaging.

### Quantitative imaging

The fully-automated ImageXpress Micro Confocal Imaging System (Molecular Devices) was used to capture both widefield and confocal images with a sCMOS camera. Widefield images were captured by a 20X Ph1S Plan Fluor ELWD ADM 0.45 NA objective. Confocal images were taken by a 40X APO LWD 1.15 NA water immersion objective and a 60-micron pinhole spinning disk. Confocal image analysis was conducted on the z-maximum projections produced from a z-series of 25 images at 0.2 micron step-sizes. Image analysis was solely performed using CellProfiler (Broad Institute)^52^. DAPI was used to mask the nucleus, and the intensities were measured within the mask. DNA content was determined using the integrated intensity of DAPI. Background levels for each fluorescence marker were determined and subtracted from the mean intensity of the given fluorescent marker in each cell. Identification and quantification of foci (γH2AX) was performed with the previously described *in-house* CellProfiler pipeline^25^.

### Nascent RNA-seq

Nascent RNA was captured and sequenced following the protocol published by Palozola et. al.^53^. Briefly, MCF10A cells were treated with either DMSO or AZD6738 for 8 hours. 5-Ethynyl Uridine (EU) (0.5 mM, Thermo Fisher Scientific, C10365) was added to the cells during the last 60 minutes of treatment. Total RNA was harvested and purified. Click-iT chemistry was used to biotinylate EU-labeled RNA which was then purified with MyOne Streptavidin T1 beads. Library generation of complementary DNA (cDNA) was conducted using the RNA bound to the beads. PCR libraries were sequenced using Novogene.

### Cell Viability Assay

Cells were seeded in 6 cm plates at ~150,000 cells per mL. Media was replaced with the respective treatment media after 24 hours. Media was refreshed with the respective treatment media every 24 hours. After 48 hours, cells were trypsinized, counted with a Countess, and reseeded in 6-well plates in fresh media. Cells were grown to confluency (~3-5 days) for the control (DMSO) well and then stained with 1 mL per well of 0.5% crystal violet staining solution (0.5 g crystal violet powder [Milipore Sigma C0775], 80 ml ddH2O, and 20 ml methanol) incubated for 20 min at room temperature on a bench rocker. The crystal violet solution was aspirated, and the plates were washed 2x with 1 mL of ddH2O. The plates were then air-dried for at least 24 hours on the benchtop, after which 1 mL of methanol per well was incubated in the well for 20 min at room temperature on a bench rocker. 200 uL of each sample and 200 uL of methanol, used as background, was then transferred to a transparent 96-well plate. The sample absorbances were then measured in triplicate at 570 nM utilizing the Tecan Spark® 20M plate reader. Data represent the mean and standard error of the mean.

### Lentivirus production and transductions

Second generation lentivirus was generated carrying the CDK2 biosensor using HEK293T cells. A transfection mixture was made with using the following: 6 μg of the DHB-mVenus plasmid (#136461), 6 μg of the psPAX2 plasmid (Addgene plasmid no. 12260) and 0.6 μg of the pMD2.g plasmid (Addgene, 12259) in Opti-MEM medium (Thermo Fisher Scientific, 31985062). The transfection mixture was then combined with 36 μl FuGENE 6 transfection reagent (Promega, E2691), according to the manufacturer’s instructions. The transfection mixture was added to HEK293 cells seeded 1 × 10^6^ cells per 100mm dish. After 8 hours, the medium was replaced, and cells were incubated for an additional 48 hours for virus production. The virus-containing medium was collected, and viral particles were concentrated using a Lenti-X concentrator (Clontech, 631232) according to the manufacturer’s protocol. Lentiviral transduction in MCF10A cells was performed by seeding cells onto 6-well plates at a density of 7 × 10^5^ cells per well. After 24 hours, the medium was replaced with medium containing 2X concentrated virus medium and 8 μg/ml polybrene (Millipore Sigma, TR-1003-G). After 24 hours, the virus-containing medium was replaced with normal growth medium. A monoclonal cell population was then isolated through single cell seeding and clonal growth.

### Live Cell Imaging

Cells were seeded in 96-well plates (Cellvis P96-1.5P) at 120,000 cells/mL. The ImageXpress**®** Micro Confocal High-Content Imaging System was used to image cells in a chamber with constant temperature (37ºC), CO2 concentration (5%), and humidity. Images were taken at 15 minute intervals for 8 hours prior to the addition of inhibitors. Media was then replaced with media containing DMSO (control), ATR inhibitor, MDM2 inhibitor or the combination of both. Time-lapse images were again taken at 15 minute intervals for an additional 24 hours. At the end of the 24 hours, the cells were fixed with 4% PFA and stained for immunofluorescence imaging of γH2AX and DAPI. The last time-lapse image of the 24 hour treatment was aligned to DAPI to determine the cytoplasmic and nuclear mean intensities of DHB-mVENUS. CDK2 activity was then calculated as the ratio of the cytoplasmic to nuclear intensity.

## DATA AVAILABILITY

Data and images are available upon request.

## ACKNOWLEDGEMENTS

We thank members of the Saldivar laboratory for helpful discussions on the work presented in this paper. C.S. is funded by a T32 Training Grant (T32GM142619) and the ARC Scholarship. C.O.M. and J.L. are funded by an award from the Cancer Early Detection Advanced Research Center (CEDAR). J.C.S. is funded by NIH grant R35GM147710 and the Breast Cancer Research Foundation award NXTGN-25-013. The research reported in this publication used computational infrastructure supported by the Office of Research Infrastructure Programs, Office of the Director, NIH, under award number S10OD034224. The content is solely the responsibility of the authors and does not necessarily represent the official views of the NIH.

## AUTHOR CONTRIBUTIONS

C.M.S and J.C.S. conceived the project and wrote the manuscript. C.M.S. conducted experiments, analyzed data, and prepared figures. C.O.M. conducted a critical experiment, and J.L. helped analyze data.

## DECLARATION OF INTERESTS

The authors declare no competing interests.

## Notes

### Competing Interest Statement

The authors have declared no competing interest.

