## Supplemental Figures for "G2-to-G0 cell cycle exit underlies sensitivity to ATR inhibition via the p53-p21-RB1 axis"

### SUPPLEMENTAL FIGURE INFORMATION

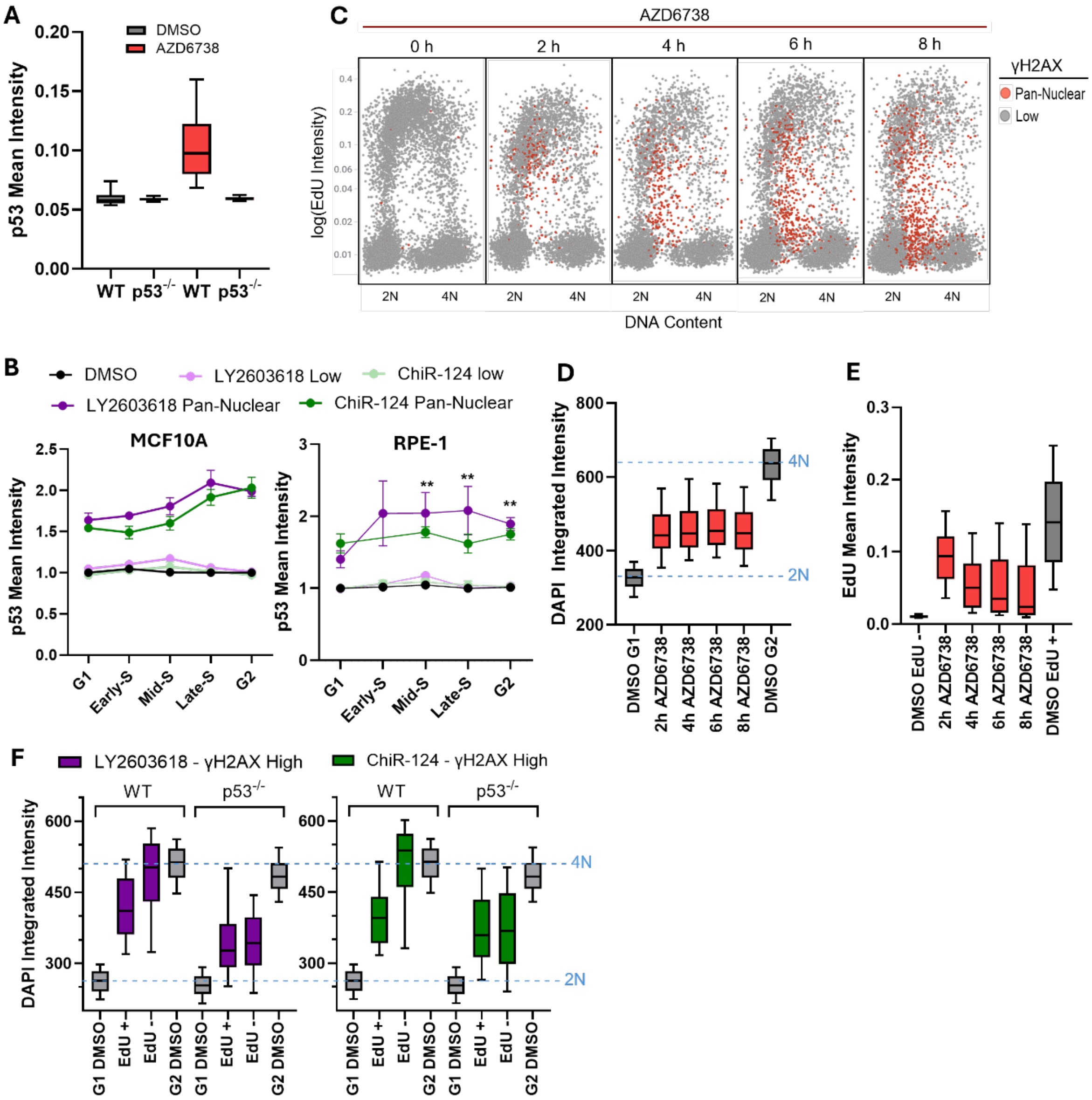


**Supplemental Figure 1. p53 is essential for DNA replication in cells with pan-nuclear γH2AX during ATR inhibition.**

A. Boxplots of p53 mean intensities in WT-MCF10A and $p53^{-/-}$ MCF10A cells during treatment with DMSO or with AZD6738 for 24 hours. Boxes and whiskers show the 25-75$^{th}$ percentile and 10-90$^{th}$ percentile, respectively.
B. Line plots of p53 mean intensity in each cell cycle phase of MCF10A and RPE-1 cells. Cells were treated as in (E).
C. Scatterplots of DNA content (DAPI integrated intensity) versus EdU mean intensity ($log_2$ scale) in $p53^{-/-}$ MCF10A treated with AZD6738 (5 µM) for 0, 2, 4, 6, or 8 hours. Cells positive for pan-nuclear γH2AX were colored red.

D. Boxplots of DAPI integrated intensities in $p53^{-/-}$ MCF10A cells treated with DMSO or 5 µM AZD6738 pan-nuclear cells (red). Boxes and whiskers show the 25-75$^{th}$ percentile and 10-90$^{th}$ percentile, respectively.
E. Boxplots of EdU Mean intensities in $p53^{-/-}$ MCF10A cells during DMSO or 5 µM AZD6738 pan-nuclear cells (red). Boxes and whiskers show the 25-75$^{th}$ percentile and 10-90$^{th}$ percentile, respectively.
F. Boxplots of DAPI integrated intensity of WT MCF10A and MCF10A $P53^{-/-}$ during treatment with DMSO (grey), 2 µM LY2603618, or 250 nM ChiR-124 for 24 hours in pan-nuclear (purple or green) damaged cells that are EdU positive or negative. Boxes and whiskers show the 25-75$^{th}$ percentile and 10-90$^{th}$ percentile, respectively.

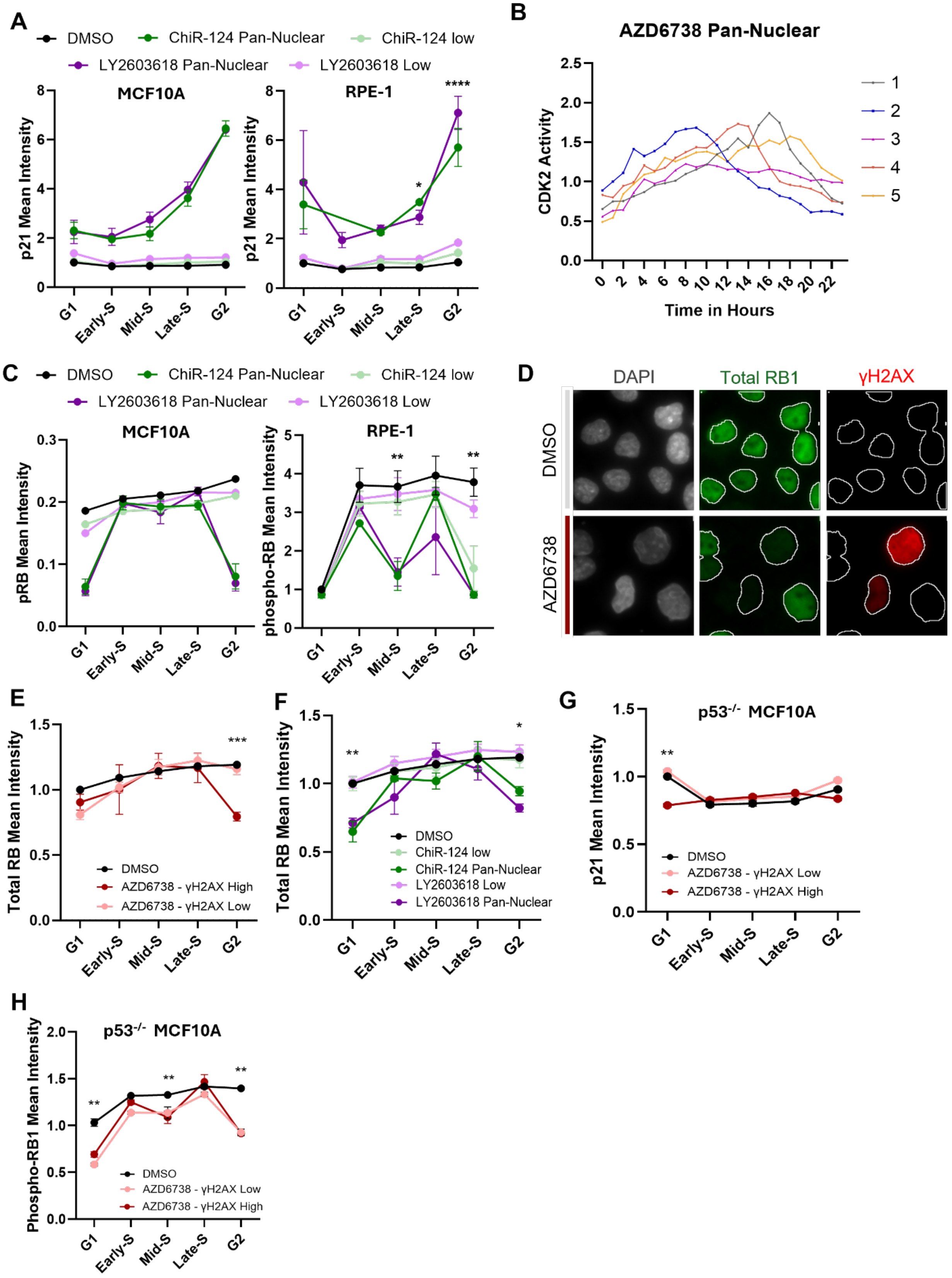


**Supplemental Figure 2. CHK1 inhibition promotes a G2-to-G0 cell cycle exit.**

A. Line plots of p21 mean intensity across the cell cycle of MCF10A and RPE-1 cells. Cells were treated with DMSO, 250 nM ChiR-124, and 2 µM LY2603618 for 24 hours.
B. Single cell traces of CDK2 activity in MCF10A-DHB cells over 24 hours of time-lapse imaging during treatment with 5 µM AZD6738.
C. Line plots of phospho-RB1 mean intensity across the cell cycle of MCF10A and RPE-1 cells. Cells were treated as in (A).
D. Representative images of DAPI, total RB1, and γH2AX. Cells were treated with DMSO or 5 µM AZD6738 for 24 hours.
E. Line plots of total RB1 mean intensity across of the cell cycle of MCF10A cells. Cells were treated as in (D).
F. Line plots of total RB1 mean intensity across the cell cycle of MCF10A cells. Cells were treated as in (A).
G. Line plots of p21 mean intensity across the cell cycle of $p53^{-/-}$ MCF10A cells. Cells were treated with DMSO or 5 µM AZD6738 for 24 hours. Dark red line shows levels in pan-nuclear γH2AX (γH2AX High) cells.
H. Line plots of phospho-RB1 mean intensity in each cell cycle phase of $p53^{-/-}$ MCF10A cells. Cells were treated as in (G). Dark red line shows levels in pan-nuclear γH2AX (γH2AX High) cells.

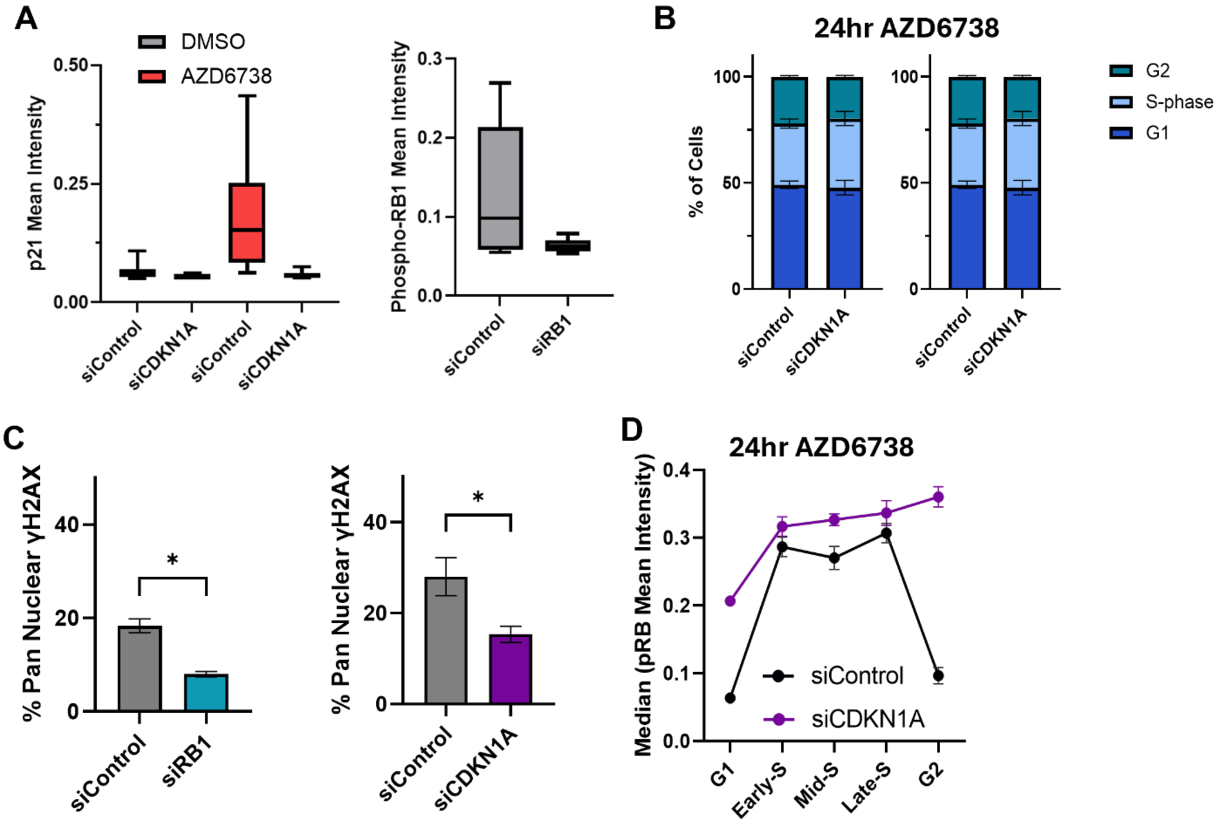


**Supplemental Figure 3. p21 and RB1 promote pan-nuclear DNA damage signaling.**

A. Boxplots of p21 and phospho-RB1 mean intensities after transfection with the respective siRNA. Cells were treated with DMSO or 5 µM AZD6738 for 24 hours. Boxes and whiskers show the 25-75th percentile and 10-90th percentile, respectively.

B. Stacked bar graphs of the percentage of cells in G0/G1, S-phase, and G2 after transfection with either siControl, siCDKN1A, or siRB1 and treated with 5 µM AZD6738.
C. Bar graphs of the percentage of pan-nuclear γH2AX cells after siRNA transfection with siControl, siCDKN1A, or siRB1.
D. Line plots of phospho-RB1 mean intensity across the cell cycle of MCF10A cells after transfection with siControl or siCDKN1A.

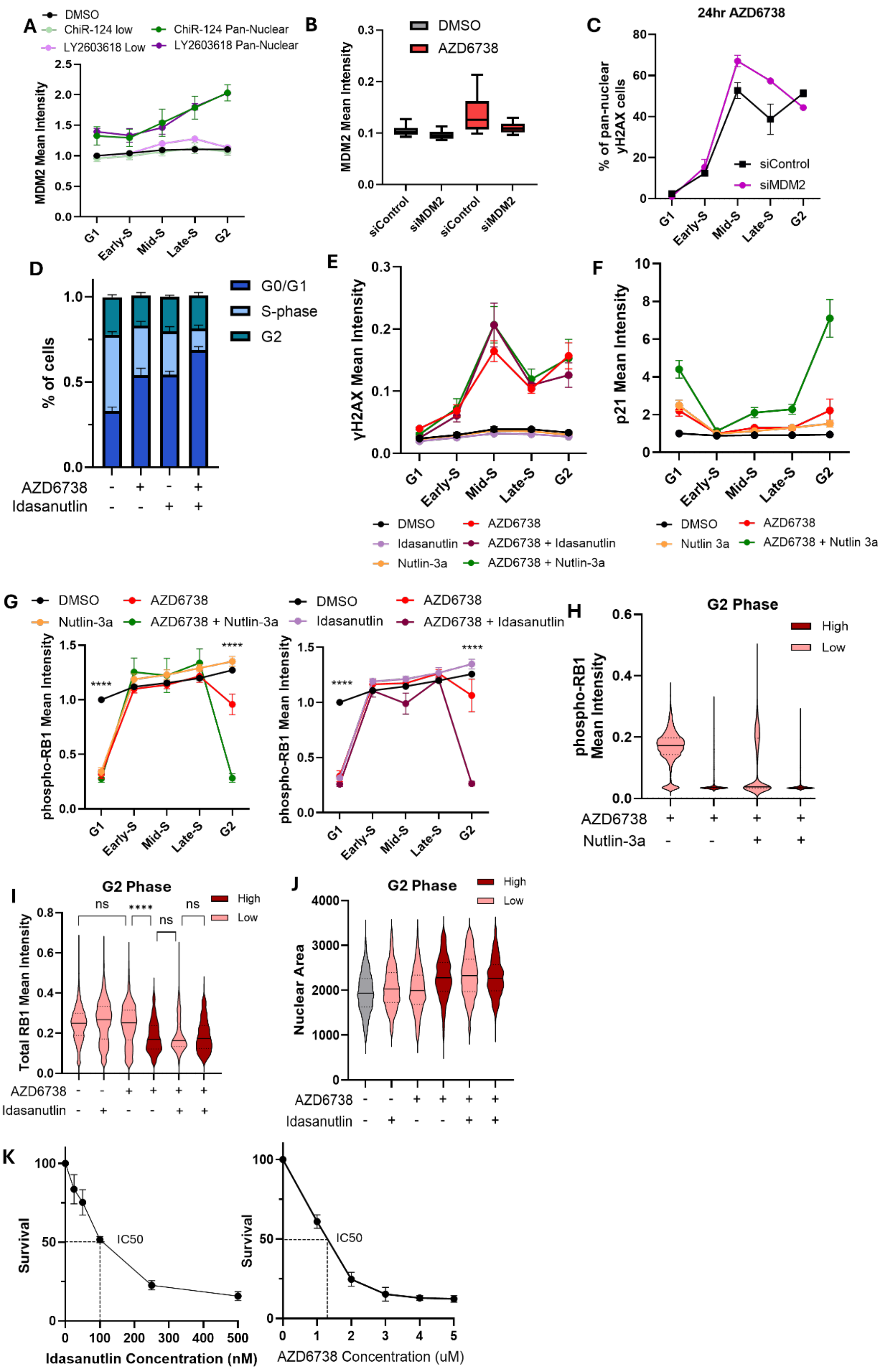

A
DMSO
ChiR-124 low
ChiR-124 Pan-Nuclear
LY2603618 Low
LY2603618 Pan-Nuclear
MDM2 Mean Intensity
G1
Early-S
Mid-S
Late-S
G2
B
DMSO
AZD6738
MDM2 Mean Intensity
siControl
siMDM2
siControl
siMDM2
C
24hr AZD6738
% of pan-nuclear yH2AX cells
siControl
siMDM2
D
G0/G1
S-phase
G2
% of cells
AZD6738
Idasanutlin
E
yH2AX Mean Intensity
DMSO
AZD6738
Idasanutlin
AZD6738 + Idasanutlin
Nutlin-3a
AZD6738 + Nutlin-3a
F
p21 Mean Intensity
DMSO
AZD6738
Nutlin 3a
AZD6738 + Nutlin 3a
G
DMSO
AZD6738
Nutlin-3a
AZD6738 + Nutlin-3a
DMSO
AZD6738
Idasanutlin
AZD6738 + Idasanutlin
phospho-RB1 Mean Intensity
****
H
G2 Phase
High
Low
phospho-RB1 Mean Intensity
AZD6738
Nutlin-3a
I
G2 Phase
High
Low
ns
****
ns
ns
Total RB1 Mean Intensity
AZD6738
Idasanutlin
J
G2 Phase
High
Low
Nuclear Area
AZD6738
Idasanutlin
K
Survival
IC50
Idasanutlin Concentration (nM)
Survival
IC50
AZD6738 Concentration (uM)

**Supplemental Figure 4. Combined MDM2 and ATR inhibition increases the G2-to-G0 cell cycle exit.**

A. Line plots of MDM2 mean intensity across the cell cycle of MCF10A cells. Cells were treated with DMSO, 250 nM ChiR-124, and 2 µM LY2603618 for 24 hours. Dark green and dark purple line shows levels in pan-nuclear γH2AX (γH2AX High) cells.
B. Boxplots of MDM2 mean intensities after transfection with the MDM2 siRNA. Cells were treated with DMSO or 5 µM AZD6738 for 24 hours. Boxes and whiskers show the 25-75$^{th}$ percentile and 10-90$^{th}$ percentile, respectively.
C. Line plots of the percentage of pan-nuclear γH2AX cells in each cell cycle phase after siRNA transfection with siControl and siMDM2. Cells were treated as in (B).
D. Stacked bar graphs of the percentage of cells in G0/G1, S-phase, and G2 after 24 hour treatment with DMSO, 50nM idasanutlin, 5 µM AZD6738, or both for 24 hours.
E. Line plots of γH2AX mean intensity across the cell cycle of MCF10A cells treated with DMSO, 50 nM idasanutlin, 1 µM nutlin-3a, 5 µM AZD6738 or the combination for 24 hours.
F. Line plots of p21 mean intensity across the cell cycle in MCF10A cells treated with DMSO, 1 µM nutlin-3a, 5 µM AZD6738, or both for 24 hours.
G. Line plots of phospho-RB1 mean intensity across the cell cycle in MCF10A treated as in (E).
H. Violin plots of phospho-RB1 intensity in the G2 phase during treatment with 5 µM AZD6738 or in combination with nutlin-3a for 24 hours. Dark red plots show levels in pan-nuclear γH2AX (γH2AX High) cells.
I. Violin plots of total RB1 intensity in the G2 phase during treatment with DMSO, 50 nM idasanutlin, 5 µM AZD6738 or the combination for 24 hours. Dark red plots show levels in pan-nuclear γH2AX (γH2AX High) cells.
J. Violin plots of nuclear area in the G2 phase. Cells were treated as in (I). Dark red plots show levels in pan-nuclear γH2AX (γH2AX High) cells.
K. Dose-response curves of MCF10A cell survival after treatment with idasanutlin and AZD6738 at the indicated concentrations.
